# Transient silencing of spinal nociceptive neurons leads to a long-term reduction in joint pain mediated by recruitment of opioid activity

**DOI:** 10.64898/2026.01.14.699276

**Authors:** Silvia Silva Hucha, Charlotte Leese, Sara Hestehave, Sara Caxaria, Sandrine M Géranton, Shafaq Sikandar, Bazbek Davletov, Stephen P Hunt, Maria Maiarú

## Abstract

Chronic inflammatory joint pain remains inadequately treated despite current therapies, creating urgent clinical need for non-addictive alternatives in an era of heightened opioid-related concerns. A promising new approach involves conjugates of botulinum toxin and substance P (SP-BOT), which, when infused locally or spinally, have been shown to produce temporary pain relief in neuropathic mouse models, with its effects declining after approximately 100 days as the construct loses activity. However, its long-term efficacy in inflammatory pain models was unknown.

Unexpectedly, in a mouse model of ankle inflammatory arthritis, a single intrathecal injection of SP-BOT at the peak of pain sensitivity produced a persistent reduction in mechanical hyperalgesia that lasted for weeks beyond the loss of construct activity. Furthermore, this late-phase pain relief exhibited a striking mechanism in that a single injection of the opioid antagonist naltrexone partially reversed the anti-hyperalgesia generated by SP-BOT.

These data indicate that SP-BOT-mediated neuronal silencing initiates two distinct phases of pain relief. An initial, direct phase is followed by a second, sustained phase that maintains pain alleviation and is driven by endogenous opioid mechanisms.

## Introduction

Chronic pain, such as that associated with arthritis, remains a significant clinical challenge, with few treatments offering prolonged relief and acceptable side effects[17; 20; 34; 45; 48]. A deeper understanding of the underlying neurobiological mechanisms is critical for developing better therapies. In arthritic conditions, pain arises from a combination of joint-specific mechanisms and central nervous system sensitization. Local factors like bone remodelling, cytokines, and growth factors sensitize peripheral nociceptors, while subsequent central sensitization in the spinal dorsal horn amplifies these nociceptive signals[20; 34; 45; 48]. While sex differences in pain processing and opioid sensitivity are well-documented [31; 42] few therapeutic development programs adequately investigate sex-specific mechanisms or treatment responses, a critical gap given that 60-70% of arthritis patients are female[39].

We have previously developed a targeted therapeutic approach using a recombinant conjugate of Substance P and the botulinum neurotoxin light chain (SP-BOT)[3; 16; 24]. This construct selectively silences NK1 receptor-expressing neurons in the spinal dorsal horn. In male mice with a peripheral nerve lesion to model neuropathic pain, a single intrathecal injection of SP-BOT produced long-lasting antinociception for approximately 100 days—consistent with the toxin’s known duration of action—without causing motor deficits or apparent toxicity[27].

However, the long-term efficacy of SP-BOT in arthritic pain, where direct nerve damage is minimal, remained unexplored. Furthermore, its effects had never been evaluated in females, despite well-established sex differences in pain processing[31; 36; 42].

To address these gaps, we investigated SP-BOT in a model of ankle joint inflammation induced by complete Freund’s adjuvant (CFA) in both male and female mice. We assessed mechanical hypersensitivity, bone remodelling via CT scans, and functional deficits through gait analysis.

Unexpectedly, we found that a single intrathecal SP-BOT injection produced a sustained attenuation of mechanical hypersensitivity that persisted for up to 140 days in both sexes, far beyond the construct’s expected active period and without signs of waning. This suggested the pain relief was being maintained by a lasting reorganization of nociceptive circuits.

Strikingly, this sustained analgesia was reversed by the opioid antagonist naltrexone implicating a recruitment of endogenous opioid mechanisms.

## Material and Methods

### Experimental Design

This study was designed to evaluate the effect of SP-BOT on inflammation-induced hypersensitivity in male and female mice. The experimenter was always blind to treatment. In all experiments, mice were randomly assigned into treatment group. The numbers of mice in each group are indicated in the individual figures.

### Animals

Reporting of these Animal used for pain studies complies with the ARRIVE guidelines. all efforts were made to minimise animal suffering (UK Animal Act, 1986) and to reduce the number of mice used (3Rs). Experiments were designed to use the minimum number of animals to provide sufficient statistical power based on our previous experience with the behavioural assay.

Subjects in all experiments were adult male and female C57BL/6J mice (8-12 weeks old) purchased form Charles River (UK) and housed to acclimatise for 1 week prior to experiments. All mice were kept in their home cage in a temperature-controlled (20 ± 1°C) environment, with a light-dark cycle of 12 hours (light on at 7:30 AM Food and water were provided *ad libitum*. All experimental protocols were approved by the institutional Animal Welfare and Ethical Review Body (AWERB) and were carried out under UK Home Office Project Licence PPL PP9720547.

### Design and Purification of Botulinum Constructs

BoNT/A consists of three domains: the binding domain, the translocation domain, and the catalytic light-chain domain, a zinc metallopeptidase. The light chain and the translocation domain form a well-packed dimer, called LcTd. We used a protein stapling technique to conjugate bacterially-expresses LcTd to chemically-synthesized substance P (SP) as described[3]. Firstly, the LcTd of the botulinum type A1 strain was fused to SNAP25 (LcTd-S25) and was prepared as previously described[12]. The chemically synthesised syntaxin-SP peptide had the sequence Ac-EIIKLENSIRELHDMFMDMAMLVESQGEMIDRIEYNVEHAVDYVE-Ahx-Ahx-RPKPQQFFGLMNH2, where Ahx stands for aminohexanoic acid. Second, the protein “staple” was prepared recombinantly using the rat vesicle-associated membrane protein 2 (VAMP2) sequence (amino acids 3–84) inserted into the XhoI site of pGEX-KG. Correctly-oriented attachment of SP to LcTD was achieved by the SNARE assembly reaction. LcTd-S25, VAMP2 (3–84), and syntaxin-SP were mixed at a molar ratio of 1:1:1 in 100 mM NaCl (sodium chloride), 20 mM Hepes, and 0.4% n-octylglucoside at pH 7.4 (buffer A). Reactions were left at 20 °C for 1 hour to allow formation of the SNARE ternary complex. SDS-resistant and irreversibly assembled protein conjugates, SP-BOT, were visualised using Coomassie Blue stain in Novex NuPAGE 12% bis-tris SDS–PAGE (polyacrylamide gel electrophoresis) gels (Invitrogen) run at 4 °C in a NuPAGE MES SDS running buffer (Invitrogen). All recombinant proteins were expressed in the BL21-Gold (DE3)pLyss strain of Escherichia coli (Agilent) in pGEX-KG vectors as glutathione S-transferase C-terminal fusion proteins cleavable by thrombin. Glutathione S-transferase fusion constructs were purified by glutathione affinity chromatography and cleaved by thrombin. Synthetic peptides were made by Peptide Synthetics Ltd.

### Mouse inflammatory model: CFA-induced ankle joint inflammation

Inflammation was induced as previously described[26]. Briefly, all mice received an injection of 5ul of CFA (Sigma-Aldrich) into the left ankle joint under isoflurane anaesthesia induced in a chamber delivering 2% isoflurane combined with 100% O2 and maintained during injection via face mask. The needle entered the ankle joint from the anterior and lateral posterior position, with the ankle held in plantar flexion to open the joint[26].

Intrathecal injections were performed under isoflurane anaesthesia[13; 27]. Briefly, the mice were held firmly by the pelvic girdle using thumb and forefinger of the nondominant hand. The skin above the iliac crest was pulled tautly to create horizontal plane where the needle was inserted. Using the other hand, the experimenter traces the spinal column of the mouse curving the column slightly to open the spaces between vertebrae. A 30-gauge needle connected to a 10μl Hamilton syringe was used to enter between the vertebrae and removed after injection. All intrathecally delivered drugs and vehicle solutions (saline) were injected in a 3μl volume.

### Drugs

Naltrexone hydrochloride (Sigma N3136) was diluted in saline and injected through the intraperitoneal (IP) route at a dose of 10mg/kg.

### Behavioural testing

All behavioural experiments were performed by experienced female experimenters, who were always blind to the treatment group; the test order was randomised with multiple groups being represented in each cage.

### Von Frey filament Test

Mice were placed in Plexiglas chambers, located on an elevated wire grid, and allowed to habituate for at least 1 hour. After this time, the plantar surface of the paw was stimulated with a series of calibrated von Frey’s monofilaments. The threshold was determined by using the up-down method[8]. The data are expressed as a log of the mean of the 50% pain threshold ± SEM.

### Catwalk gait analysis

Analysis of voluntary movement and gait pattern was performed using the Catwalk XT 10.0 system (Noldus Information Technology) [49], and based on our previous experience[21]. Briefly, green light was internally reflected into a glass plate, on which an enclosed corridor was fixed, with red backlight above the corridor. A video-camera was mounted underneath the setup and recorded the paw prints being lit up by the green light when paws were in contact with the glass plate as the animal walked along the corridor. A run was regarded as compliant when the animal entered in one end of the corridor and moved fluently across the plate towards the other end of the corridor, with a running duration below 12 seconds and a maximum variation below 75%. Three compliant runs were recorded for each animal, with no previous training/habituation or food-deprivation. Following the recording, compliant runs were classified and manually cleaned/corrected for potential miscellaneous prints. For outcome-measures like mean intensity, the data was converted into a ratio between ipsi- and contra-lateral hind-limbs: (LH/RH) / 100%. The parameter “guarding index” was calculated as previously described[1], and a higher guarding index suggests less dynamic weight bearing on the injured leg compared with the non-injured leg.

### Ankle swelling

Ankle swelling was assessed using a precision millimetric calliper, following a standardised protocol to ensure consistency and reproducibility. Measurement was taken at the end of all behavioural experiments. Each animal’s ankle was measured from two orthogonal planes: the anterior-posterior (frontal) view and the medial-lateral view. The calliper was gently positioned to avoid compressing the tissue, and care was taken to maintain the same anatomical landmarks across subjects and time points.

The two linear measurements obtained from each mouse were used to estimate the ankle perimeter by applying an elliptical approximation formula[4], allowing for a more accurate quantification of soft tissue swelling. This method was applied to animals treated with the SP-BOT conjugate as well as those receiving vehicle control (saline solution). Comparative analysis of the calculated ankle perimeters enabled the evaluation of inflammatory oedema across treatment groups and between sexes. Statistical comparisons were performed to determine whether SP-BOT treatment differentially modulated CFA-induced inflammation in male and female mice.

### mCT imaging and analysis

Ankles were imaged for microCT analyses using VECTor6CT (MILabs, The Netherlands) with the following settings: ultra-focus magnification, accurate scan mode, default settings, for a total duration of approximately 5min. Reconstruction was done using 20voxel size. Images were analysed using VivoQuant version 4 software (Invicro LLC, Boston, MA, USA) as follows: ROIs selected for end of tibia, calcaneus and talus and bone volume quantified. Bone volume for each mouse was done for ipsilateral and contralateral ankle and a ratio for the volume calculated as ipsilateral/contralateral volume[22].

### Immunohistochemistry

Mice were anaesthetised with pentobarbital and perfused transcardially with physiological saline containing heparin (5,000 IU/mL), followed by 4% paraformaldehyde (PFA) in a 0.1 M phosphate buffer (PB; 25 mL per adult mouse). Joints were collected from some mice for microCT scanning and fixed in PFA for at least 7d. Lumbar spinal cords were dissected out, fixed in 4% PFA for an additional 2 hours, and transferred into a 30% sucrose solution in a PB containing 0.01% azide at 4 °C for a minimum of 24 hours. Spinal cord sections were cut on a freezing microtome set at 40 µm. Sections were left to incubate with the primary antibodies (rabbit-NK1R [Sigma], rabbit vesicular glutamate transporter 2 (VGLUT2) [Synaptic System], rabbit potassium-chloride cotransporter 2 (KCC2) [Millipore], guinea pig- vesicular GABA transporter (VGAT) [Synaptic System], guinea pig- calcitonin gene-related peptide (CGRP) [Chemicon]) and rabbit anti-cSNAP25 recognizing the cleaved end of SNAP25, TRIDEANQ 1:50,000[28])

Biotinylated secondary antibody (biotinylated secondary antibody goat anti-rabbit, Vector Laboratories BA-1000) was used for NK1 staining at the concentration of 1:400 in PBS-T solution and left for 90 minutes at room temperature. Then, sections were incubated with avidin-biotin complex in PBS-T (1:250 Vectastain A plus and 1:250 Vectastain B; ABC Kit Peroxidase Standard, Vector Laboratories) for 30 minutes at room temperature, followed by a signal amplification step with biotinylated tyramide solution (1:75 for 5 minutes; Startech). Finally, sections were incubated with Fluorescein Avidin D (1:600, Vector Laboratories) for 2 hours at room temperature. For all the other staining, appropriate fluorescent secondary antibodies (1:500, Jackson Research) were used. All fluorescent sections were transferred to glass slides and cover slips applied with Gel Mount Aqueous Mounting Medium (Sigma-Aldrich) to prevent fading and stored in dark boxes at 4 °C.

### Quantification of Fluorescence

Quantitative analysis of optical density (OD) was performed using ImageJ software (NIH, USA). OD measurements were obtained specifically over laminae I–II of the superficial dorsal horn in the spinal cord, focusing on regions exhibiting immunofluorescent labelling. For each region of interest (ROI), the mean value, representing the OD, was recorded. To correct for non-specific background fluorescence, background OD values were acquired from adjacent areas within the same tissue section that lacked specific labelling. A total of four sections per animal were analyzed, ensuring representative sampling. Contrast enhancement and fluorescence threshold were kept constant. The resulting OD values were averaged per animal and used for subsequent statistical comparisons.

### Data and Statistical Analysis

All experimental data points were included in the data analysis. All statistical tests were performed using the IBM SPSS Statistic programme (version 29), and P < 0.05 was considered statistically significant. For von Frey data, statistical analysis was performed on the data normalised by log transformation. Please note that, as in our previous paper[26; 27], we logged the data of von Frey filaments test to ensure a normal distribution because the von Frey’s hairs are distributed on an exponential scale. Mills et al [30] demonstrated that log transformation makes more “mathematical and biological sense”. Difference in sensitivity was assessed using repeated measure mixed model analysis of Variance (ANOVA). In all cases, “time” was treated as a within-subjects factor and “treatment” was treated as a between-subject factor.

For gait analysis, each sex was tested in independent cohorts, the statistical analysis is conducted independently per sex, using unpaired one-tailed t-tests for comparison between vehicle and SP-BOT treated groups.

For immunofluorescence quantification, the results were normalised to the mean of the control-group (contralateral side of the saline treated mice), and the control mean was set as 100%.

## Results

Building on previous work demonstrating that a single intrathecal injection of SP-BOT can transiently silence NK1R-positive neurons and reduce hypersensitivity in male mice[26], and more recent evidence of its long-term protective effects against neuropathic pain for around 100 days [27] we investigated its long-term efficacy in a model of arthritis in both male and female mice. We assessed mechanical allodynia, gait dynamics, and bone remodelling following a single intrathecal injection of SP-BOT administered after the full development of CFA-induced hypersensitivity.

### SP-BOT produces a sustained reduction in mechanical hypersensitivity

Previous research in naïve mice has shown that intrathecal SP-BOT administration produces no motor impairment or changes in baseline mechanical thresholds[26]. However, when administered on day 7 after CFA injection, it produced a substantial and sustained reduction in mechanical hypersensitivity in both male and female mice, an effect that persisted throughout the experiment and remained statistically significant at day 128 (Fig. 1B and D). As expected, hypersensitivity was not observed in the contralateral paw (Fig. S1), and SP-BOT treatment did not affect CFA-induced ankle swelling (Fig. 1C and E). This dissociation between analgesia and persistent peripheral inflammation is notable and indicates that pain relief operates independently of anti-inflammatory mechanisms.

**Figure 1.**
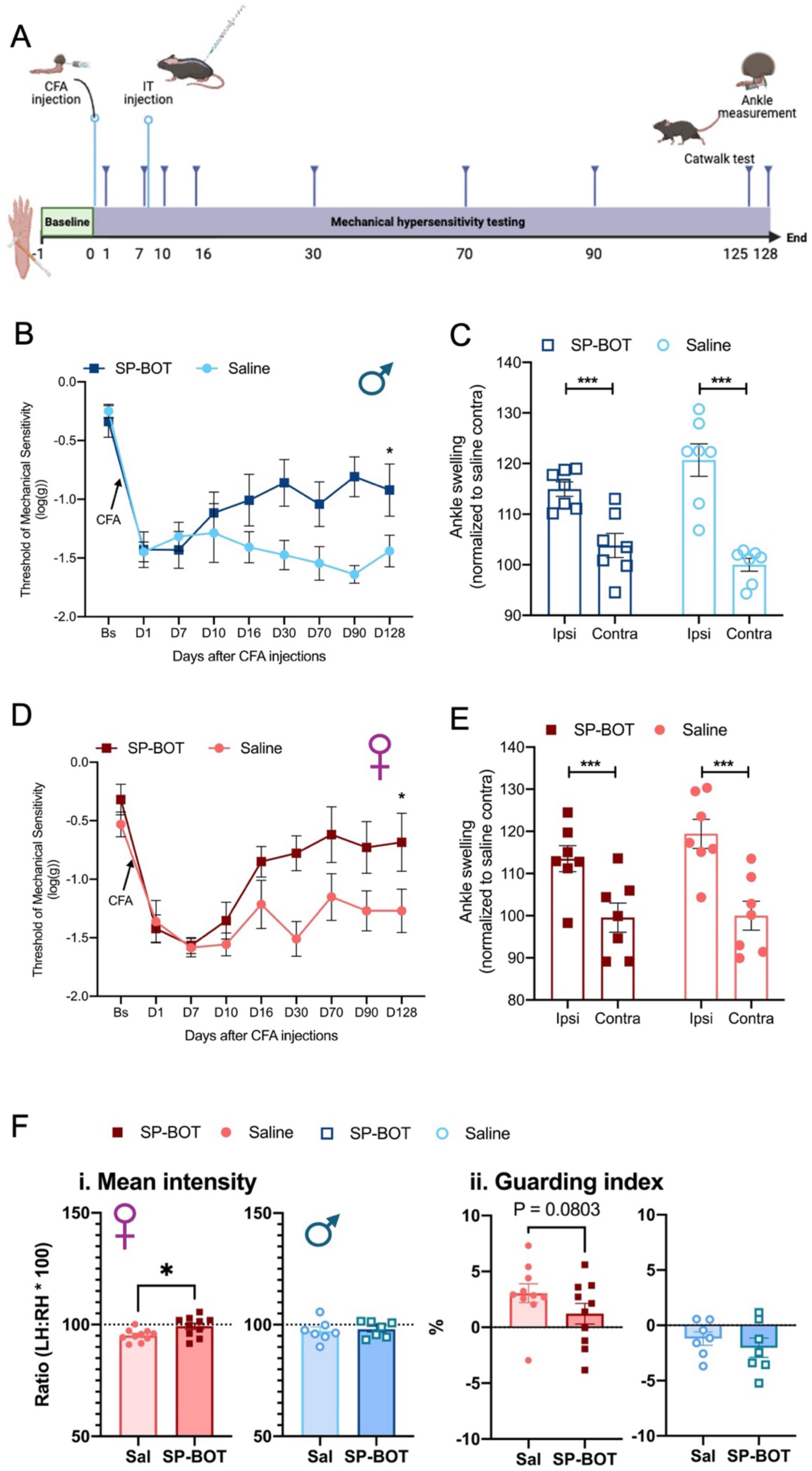
SP-BOT attenuated CFA-induced mechanical hypersensitivity in male and female mice and restored weight-bearing and gait patterns in females. **(A)** Schematic of experiment timeline. Von Frey test was performed before and after CFA injection and tracked out for 128 days. SP-BOT (100ng/ 3ul) was administered intrathecally 7 days post-CFA, when mechanical hypersensitivity had developed. (B and D)) Mechanical sensitivity thresholds (log(g)) over time following CFA injection (5ul) into the ankle joint (black arrows) in male **(B)** and female **(D)** mice. SP-BOT treatment significantly alleviated mechanical hypersensitivity in both sexes compared to saline controls. Mice were tested at baseline and up to 128 days after CFA injection. Difference in sensitivity was assessed using repeated measure mixed model two-way ANOVA (male, n=8/8, F_1,14_ = 10.5, P = 0.006, Bonferroni correction at D128, P=0.016; female, n=8/8, F_1,14_ = 5.7, P = 0.031, Bonferroni correction at D128, P=0.02). **(C, E)** Ankle swelling normalized to contralateral control ankle at D128 in males **(C)** and females **(E)**. Both SP-BOT and saline-treated animals show significant ipsilateral swelling (***p < 0.001), with no significant difference between treatment groups. Male, Two-way ANOVA, factor Side, F = 51.8, P < 0.001; factor Treatment, F = 0.19, P = 0.67. Female, Two-way ANOVA, factor Side, F = 21.5, P < 0.001; factor Treatment, F = 1.05, P = 0.33. Data are presented as mean ± SEM. *p < 0.05, ***p < 0.001. **(F)** Gait deficits were recorded using the Catwalk XT system, recording both female (closed symbols) and male (open symbols) mice 125 days after CFA injection into the left ankle joint. SP-BOT was administered at day 7 after injury. **i)** Mean intensity of the left hind-paw print was decreased by CFA-injection (vehicle) and normalized by SP-BOT treatment for females, but unaltered for males. **ii)** The guarding of the left leg in females was increased by CFA-injury (vehicle-treated) but partially restored by SP-BOT (trend; P= 0.0803). No significant guarding was seen in male mice, and the guarding index was unaltered by treatment. Males: N=7, females: N=10 per treatment-group. Error bars suggest mean ± S.E.M. Statistical comparison between vehicle and SP-BOT treated groups were conducted using unpaired one-sided t-tests, and significant results were marked by: *P<0.05.

### SP-BOT rescues late-stage gait deficits in female mice

To objectively assess long-term functional benefits, we analyzed gait patterns at 125 days post-CFA injection. At this late stage, CFA-induced gait alterations in male mice were mild and unaffected by SP-BOT treatment (Fig. 1Fi, Fii). In female mice, however, CFA injection induced significant deficits, seen as decreased mean intensity and an increased guarding index, suggesting reduced dynamic weight-bearing on the injured leg. Early treatment with SP-BOT restored these parameters to near-normal levels (Fig. 1Fi, Fii), indicating a long-lasting functional improvement.

### SP-BOT does not alter CFA-induced bone pathology

CFA injection induced significant pathological bone remodelling, evidenced by a marked increase in ipsilateral bone volume (Fig. 2A, B). Intrathecal SP-BOT administration had no detectable effect on this bone pathology in either sex (Fig. 2C, D), confirming that its potent anti-nociceptive and functional benefits occur independently of changes in the underlying joint structure. Additionally, this also suggests that despite a potential increased load on the injured leg, due to the analgesic effects of SP-BOT, this did not worsen the pathological condition.

**Figure 2:**
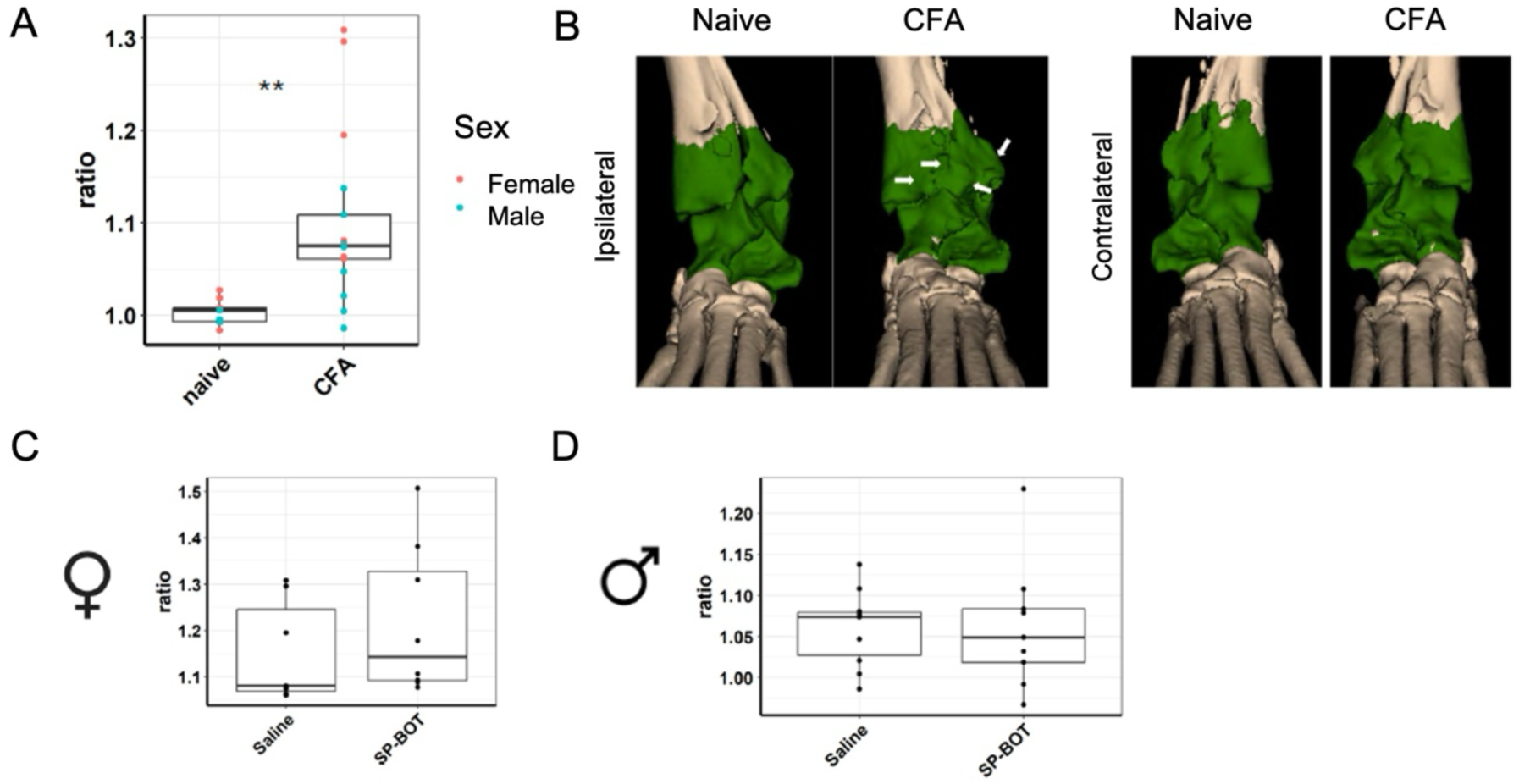
Persistent osteophyte formation and bone volume changes following CFA injection are not attenuated by intrathecal SP-BOT. **(A)** Bone volume ratio for naïve and CFA injected mice as ratio between ipsilateral and contralateral ankles of the hind paws. Naïve mice have ratio ∼1 indicating similar ankle volumes between legs while CFA injected ankles show an increase in bone volume following CFA injection. Animals colour coordinated by sex. Results were compared using Student t-test (naïve n=10, CFA =17) * for p< 0.05, ** for p<0.01 **(B)** Representative images for ipsilateral and contralateral ankles for naïve and CFA injected male mice. White arrows indicate bone changes and osteophytes which result in increased ratio as seen in A. (**C-D**) Bone volume ratio for CFA injected mice treated with intrathecal injection of vehicle or SP-BOT (100ng/3μl) showing no changes in bone volume for both male and female mice. Results were compared using Student t-test (C: saline n=7 SP-BOT n=8; D saline n=10 SP-BOT n=9) * for p< 0.05, ** for p<0.01.

### Immunohistochemical analysis after SP-BOT

To determine if the sustained behavioural effects of SP-BOT were accompanied by structural or molecular changes in the spinal cord, we first confirmed that its mechanism of action remains reversible. Consistent with its design, SP-BOT-induced silencing did not cause ablation of NK1R+ neurons (**Fig 3 A-B**) and cSNAP immunoreactivity (a marker of SP-BOT construct activity) was almost completely lost from the dorsal horn of SP-BOT treated mice at 90d and 120d (**Fig S2)**. Analysis 120 days post-injection revealed no difference in NK1R-immunoreactivity in the superficial dorsal horn between SP-BOT and saline-treated mice of either sex (**Fig. 3 C-D**), confirming the absence of long-term neuronal loss.

**Figure 3:**
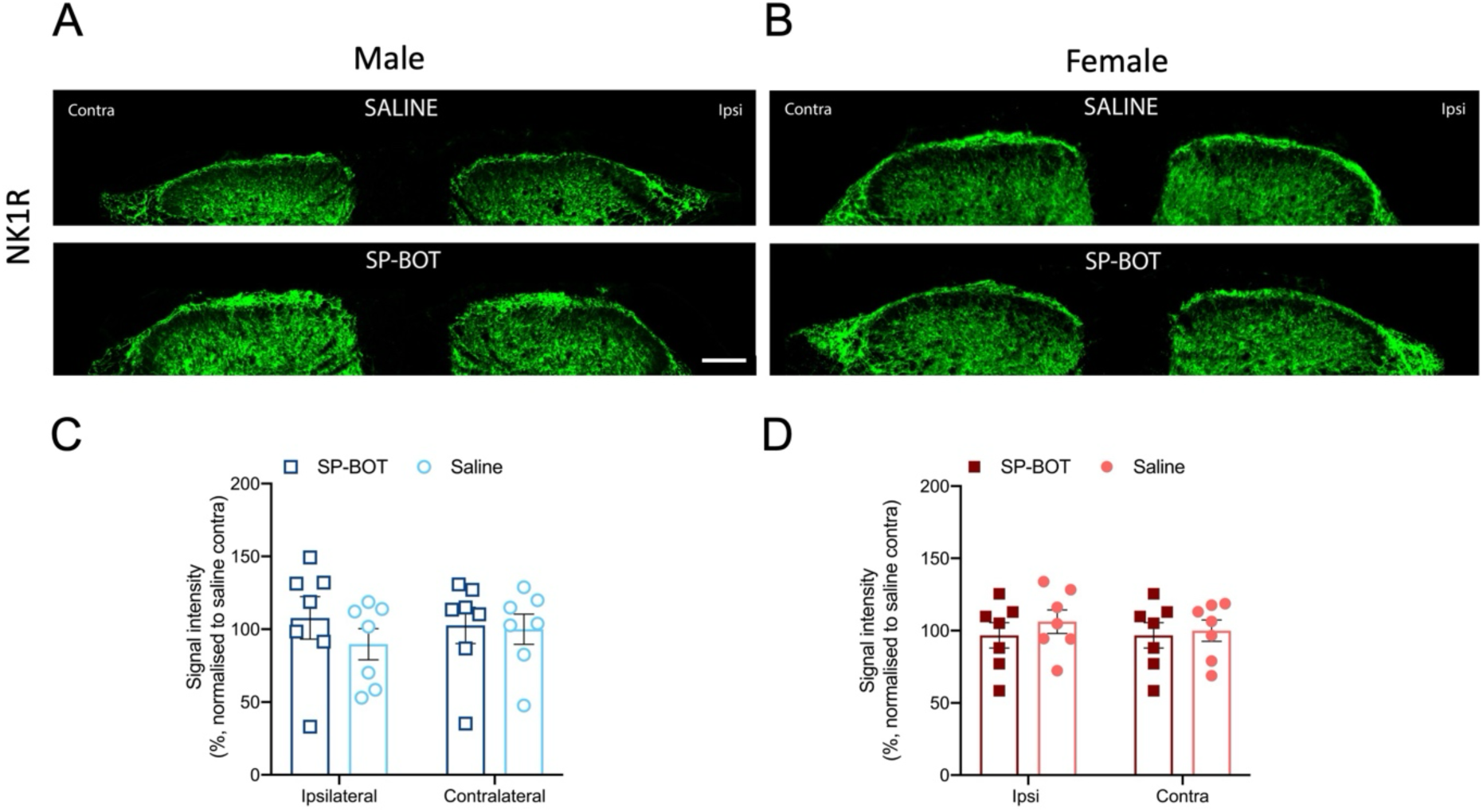
Intrathecal injection of SP-BOT did not reduce NK1-receptor immunoreactivity in the superficial dorsal horn. **(A-B)** Representative images of NK1R immunoreactivity in the superficial dorsal horn of male and female mice 130 days after injection of CFA in the left ankle joint and 123 days after intrathecal injection of SP-BOT or saline vehicle. Green, NK1R. Scale bar, 100μm. **(C-D)** quantification of NK1R fluorescence in the ipsilateral and contralateral superficial dorsal horn of male and female mice 130 days after CFA injection and 123 days after intrathecal injection of SP-BOT or saline vehicle. All data were normalised to contralateral saline-treated mice (n=7 per group). Bars represent mean ± SEM on the average fluorescent intensity.

We next analysed whether the prolonged silencing of NK1R+ neurons induced compensatory changes in key proteins involved in spinal pain processing. In male mice, we observed a bilateral increase in VGLUT2 immunoreactivity in the superficial dorsal horn following SP-BOT treatment (**Fig. 4A and C**), suggesting a potential adaptive increase in excitatory synaptic machinery. This change was not observed in female mice (**Fig. 4B and D**). Given that short-term (4w) chronic inflammation can disrupt spinal inhibition[25], we also assessed markers of inhibitory signalling. However, in contrast to previous use of shorter survival times[25], we detected no significant differences between ipsilateral and contralateral sides in the intensity of CGRP, KCC2, or VGAT signals in either sex (**Fig S3, S4, S5**). This implies that the potent and sustained anti-nociceptive effects that follow SP-BOT treatment are not mediated by preventing a CFA-induced loss of spinal inhibition, nor do they cause a widespread dysregulation of the key neurochemical markers we assessed.

**Figure 4:**
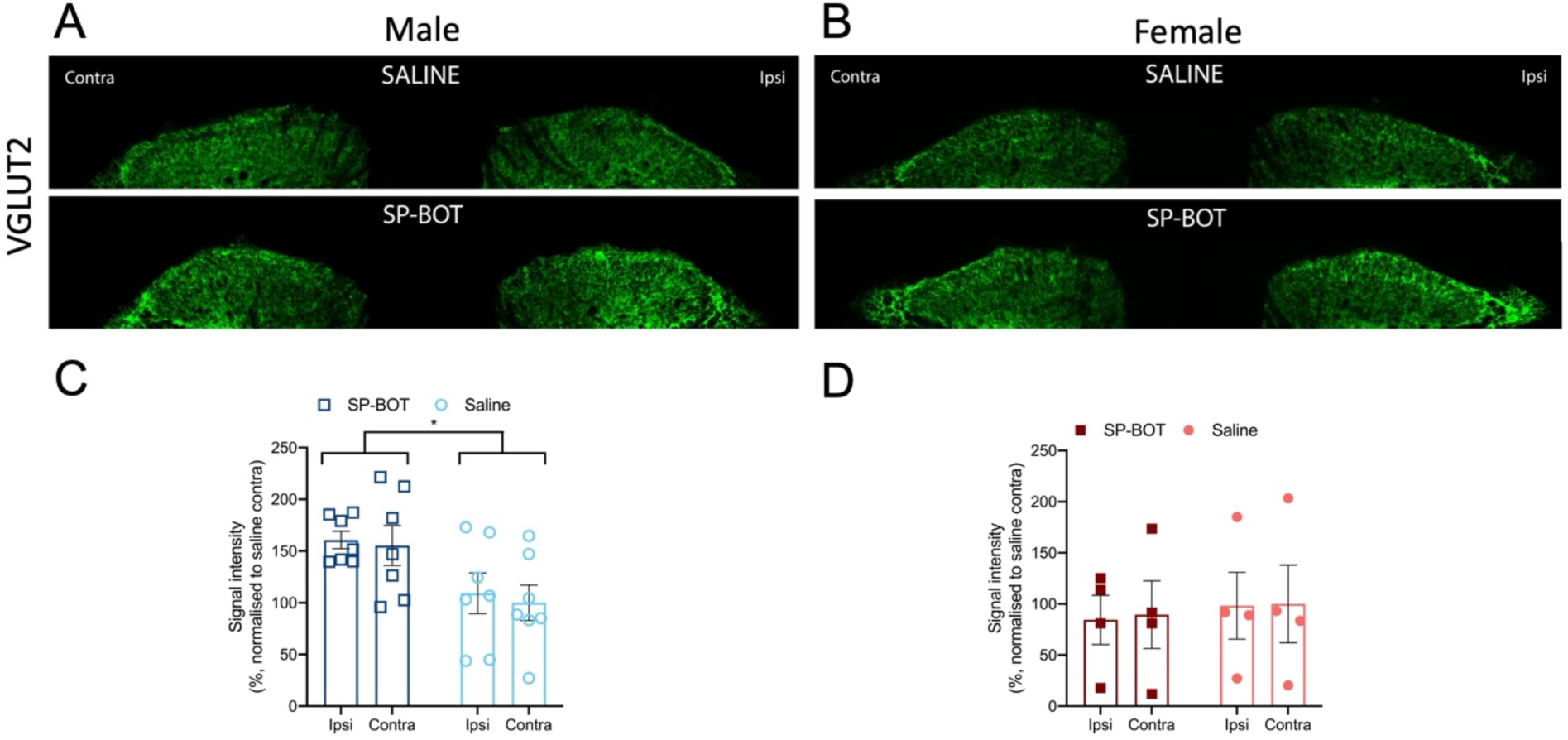
Intrathecal injection of SP-BOT increases VGLUT2 immunoreactivity in the superficial dorsal horn of male mice. **(A-B)** Representative images of VGLUT2 immunoreactivity in the superficial dorsal horn of male and female mice 130 days after CFA injection. Scale bar, 100μm. **(C-D)** Quantification of VGLUT2 fluorescence in the ipsilateral and contralateral superficial dorsal horn of male and female mice 130 days after injection of CFA in the left ankle joint after intrathecal injection of SP-BOT or saline vehicle. All data were normalised to contralateral side of saline-treated mice (male, n=7; female, n= 4). Bars represent mean ± SEM on the average fluorescent intensity. Two-Way Anova, Male, factor Treatment, F = 6, P = 0.0302; factor side, P=0.45; female, factor Treatment, P=0.78; factor Side, P=0.84. *P<0.05

### Opioid antagonism reverses SP-BOT-mediated analgesia in female mice

To further investigate the long-term effects of SP-BOT on pain hypersensitivity and explore potential sex differences in the underlying mechanisms, male and female mice were injected with CFA into the ankle joint. Once maximal mechanical hypersensitivity had developed (Day 7), animals received a single intrathecal injection of SP-BOT. As expected, this treatment resulted in a sustained reduction of hypersensitivity in both sexes. To assess whether endogenous opioid mechanisms contributed to this prolonged analgesic effect, mice were administered an intraperitoneal injection of naltrexone hydrochloride (NTX) (10mg/kg) 150 days after CFA treatment. Strikingly, overall NTX induced a marked reinstatement of mechanical hypersensitivity (**Fig 5B-C).** While the effect was strong in females it was only evident in two of the four male mice (**Fig 5A-C).** Male animals, n=4 was due to one unrelated health event leading to the exclusion of one male mouse from day 128 onwards. The smaller male sample size may limit statistical power for sex comparisons; however, the uniform response across all female animals (n=5) demonstrates robust effects on opioid activity.

**Figure 5.**
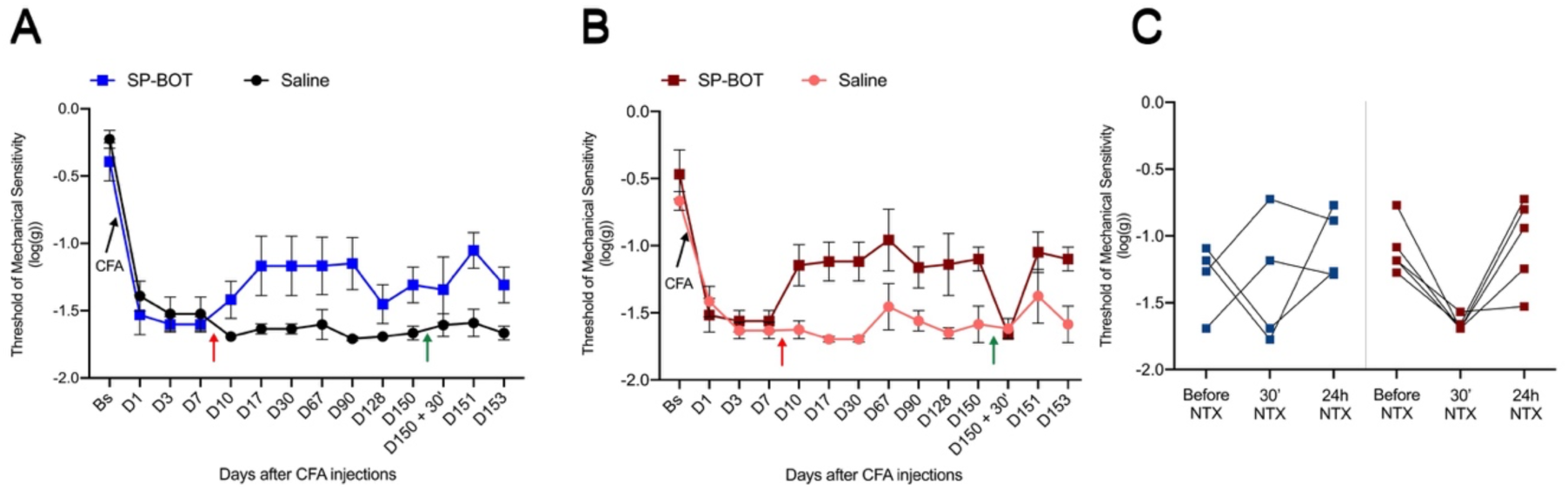
Long-lasting analgesic effect of intrathecal SP-BOT in CFA-injected male and female mice and naltrexone-induced reinstatement of hypersensitivity. Graphs show mechanical sensitivity thresholds (log(g)) over time in male (**A**) and female (**B**) mice following CFA injection into the ankle joint (black arrows, Day 0), which induced robust mechanical hypersensitivity in both sexes. On Day 7 (red arrows), mice received a single intrathecal injection of either SP-BOT or saline. SP-BOT treatment significantly attenuated hypersensitivity in both males and females for several weeks. On Day 150, all mice received an intraperitoneal injection of naltrexone (green arrows). Difference in sensitivity was assessed using repeated measure mixed model two-way ANOVA (male, Factor ‘Time x Treatment’, F=2.99, P = 0.0011; female, Factor ‘Time x Treatment’, F = 11.3, P = 0.002) (**C**) Naltrexone (10mg/kg) induced a pronounced reinstatement of hypersensitivity in female mice treated with SP-BOT but increased hypersensitivity was only seen in two of four male mice. (Repeated measure mixed model two-way ANOVA, D150 to D151, Within-Subjects, Factor TIME, F=5.37, P=0.19; Factor ‘TIME*SEX’, F=1.79, P=0,2; Between-Subjects, Factor ‘SEX’ F_1,7_= 0.11, P= 0.7). Data are presented as mean ± SEM (n = 5 per group; male SP-BOT n=4 from day 128).

## Discussion

### NK1R+ Neurons as a Therapeutic Target

NK1 receptor-expressing (NK1R+) neurons in the spinal dorsal horn are crucial for maintaining persistent pain states[23; 29; 32]. NK1R expressing neurons are found in superficial and deep laminae of the spinal cord including a key population of nociceptive neurons in lamina 1 that project to the brainstem as well as numerous interneurons in deeper laminae[7; 41]. The exact proportion of nociceptive projection neurons that express NK1R in lamina 1 is debated. Cameron et al using immunohistochemistry estimated greater than 90% [7] while others using other approaches find a smaller proportion and highlight the possibility of parallel nociceptive pathways to the hindbrain[9; 41].

NK1R positive neurons in the superficial dorsal horn of the spinal cord to the brain have been shown to be essential for the maintenance of inflammatory and neuropathic persistent pain states[32]. The original studies showed that ablation of NK1R positive neurons with intrathecal application of substance P-saporin (SP-SAP) conjugates resulted in normal acute thermal and mechanical stimulus detection but a loss of mechanical and thermal hypersensitivity in inflammatory and neuropathic pain models. Essentially, dorsal horn neurons that possess the NK1R were thought to play a pivotal role in the development of central sensitization and hyperalgesia[29; 32; 44; 47].

While these studies using the cytotoxin conjugate Substance P-saporin (SP-SAP) to ablate NK1R+ neurons validated their role and indeed translational studies in dogs also showed that intrathecal SP-SAP treatment alleviated bone cancer pain[6], our studies have employed SP-BOT, a conjugate that reversibly silences—but does not kill—NK1R+ neurons. The botulinum neurotoxin cleaves SNAP25, inhibiting synaptic release. We confirmed that activity of SP-BOT is transient, with a substantial loss of cleaved SNAP25 immunoreactivity by 90 days post-injection aligning with the known duration of botulinum toxin’s biological efficacy in a long-term model of neuropathic pain[2; 5; 35; 38]. The persistent analgesia observed in the current inflammatory pain model was therefore surprising and pointed to an indirect, long-term adaptive mechanism.

### Opioid Signalling and Latent Sensitization

The partial reversal of mechanical hyperalgesia by naltrexone points to compensatory opioid signalling. This aligns with the concept of “latent sensitization,” where a resolved pain state is actively masked by endogenous opioid tone that masks a persistent central sensitization[19]. Corder et al [10] were able to show that the mechanical hyperalgesia that accompanies acute inflammation could be reinstated by opiate receptor antagonist naltrexone [52]. Further studies also revealed that the underlying central sensitization that drives increased mechanical sensitivity following injury was still present in the dorsal horn even when the pain state had resolved but masked by increased opioid tone within the dorsal horn. This was true of both male and female mice. It was also shown that the increased opioid tone was bilateral and, in male mice, required the expression of opiate receptors on the terminals of primary afferent nociceptors within the dorsal horn. In female mice opioid receptors on dorsal horn neurons as well as primary afferent nociceptors were thought to play a role in the masking of central sensitization and hyperalgesia[40; 51].

It is important to note that in studies of latent sensitization, opioid peptide knockout mice showed recovery from hyperalgesia and reinstatement by naltrexone implying that endogenous opioid peptides were not required for an opioid receptor-mediated suppression of the hyperalgesic state but constitutive activity of the opioid receptor[10; 40; 50] was essential. However, there is evidence that descending pathways from the brainstem and local dorsal horn networks may be involved as latent sensitization to pain is modulated by the activation of 2A adrenergic receptors and Neuropeptide Y receptors[43; 50]. It has also been shown that stress is able to block the suppression of hyperalgesia during latent sensitization also implicating descending pain pathways[37].

In the present study we found very little evidence for long-term changes in inhibitory nociceptive networks within the dorsal horn[25]. Indeed, in male mice only we found a bilateral increase in VGLUT2 and which is predominantly found in excitatory dorsal horn interneurons that may also co-release enkephalin[46]. It is unclear what this might contribute to the hyperalgesic state in male mice, but it does point to different neural mechanisms in male versus female mice. Our data are incomplete on this point. While all female mice showed an increase in mechanical sensitization following naltrexone treatment, the male response was partial. This may have been due to effects of stress particularly as gender related differences have been reported or to the low number of male mice available[15].

We propose that the initial silencing of NK1R+ neurons by SP-BOT triggers a compensatory increase in opioid tone, likely mediated by opioid receptors on central neurons and nociceptive axon terminals within the dorsal horn. This could explain why the effect persists in inflammatory pain (where opioid receptors are preserved) but not beyond 100 days in SNI neuropathic pain models where nerve injury causes a loss of opioid receptors on primary afferent nociceptors[11; 33]. The incomplete naltrexone-insensitive analgesia in some SP-BOT treated male mice implies the engagement of alternative, non-opioid pathways perhaps involving descending pathways from the brainstem[14; 18; 44], and which requires further investigation.

### Conclusion

This research demonstrates the promising clinical utility of SP-BOT in treating chronic inflammatory joint pain, providing long-lasting pain relief while preserving peripheral joint structure. The research also uncovers a neuroadaptive mechanism whereby transient neuronal silencing can activate lasting endogenous pain-control pathways. Rather than providing chronic exogenous opioids, this approach activates endogenous opioid-mediated analgesia, suggesting development of therapeutics that trigger these natural pain-control mechanisms could provide non-addictive, sustained relief.

## Author Contributions

Study conception and experimental designs: *MM and SPH contributed equally to the project. Data collection: All behavioural studies were done by SSH and MM. Tissue collection and analysis by SSH, MM and SPH. Catwalk was performed by SH and SSH. SC and SS acquired and analysed joint micro-CT scans. CL and BD synthesised SP-BOT. Data analysis and interpretation: SH and MM analysed and interpreted all data. SG provided critical insights throughout the study. Manuscript preparation: SPH and MM wrote the initial drafts of the manuscript. All authors provided critical comments and revisions on the draft and all authors read and approved the final manuscript.

## Supplementary files

**Figure S1:**
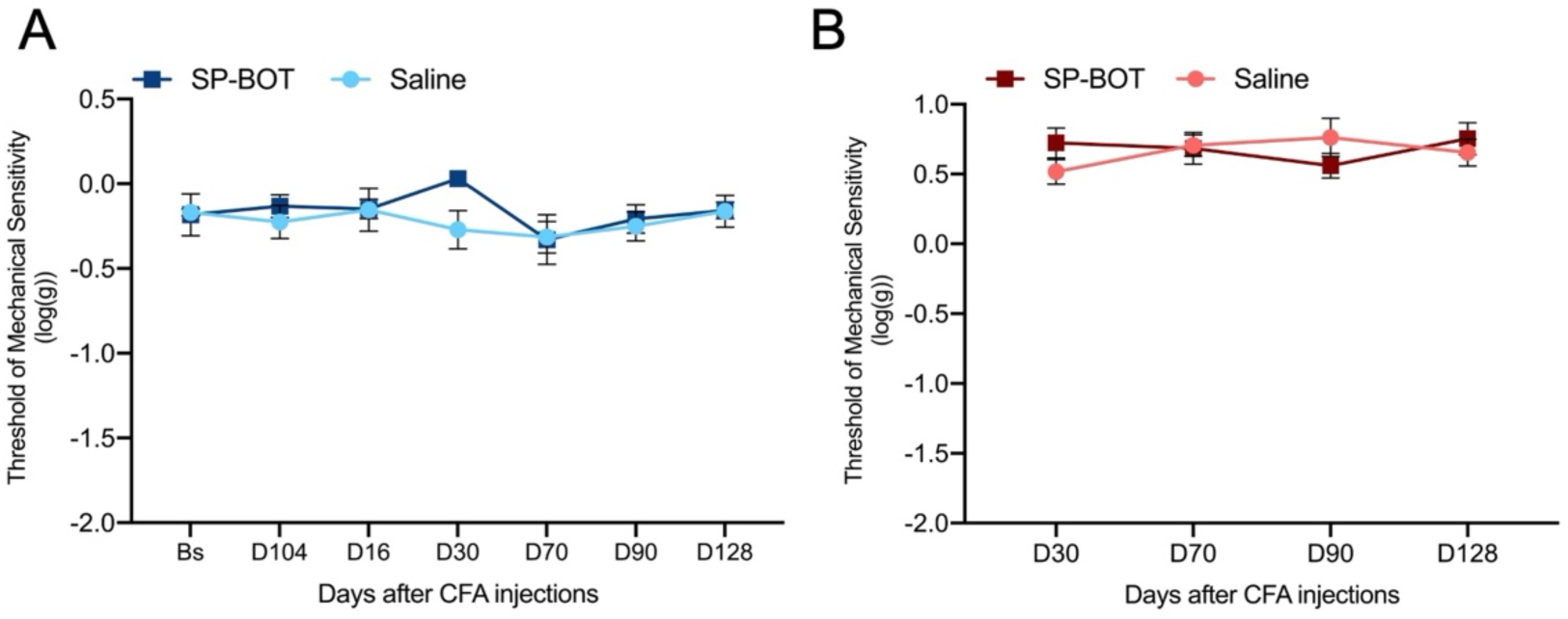
Contralateral paw mechanical sensitivity remains unaffected following ipsilateral CFA injection regardless of sex or intrathecal SP-BOT treatment. (A) Mechanical nociceptive threshold (von Frey test) of the contralateral hind paw in male mice receiving intrathecal SP-BOT or saline following CFA injection into the ipsilateral ankle joint. No significant changes in contralateral mechanical sensitivity were observed in either treatment group from baseline (Bs) through day 128 (D128), indicating that nociceptive sensitization is localized to the ipsilateral injected paw. (B) Mechanical nociceptive threshold of the contralateral hind paw in female mice. Similar to males, both SP-BOT and saline-treated females maintained normal contralateral paw sensitivity throughout the 128-day observation period, with no treatment effect observed. Data presented as mean ± SEM. (n=8/8)

**Figure S2:**
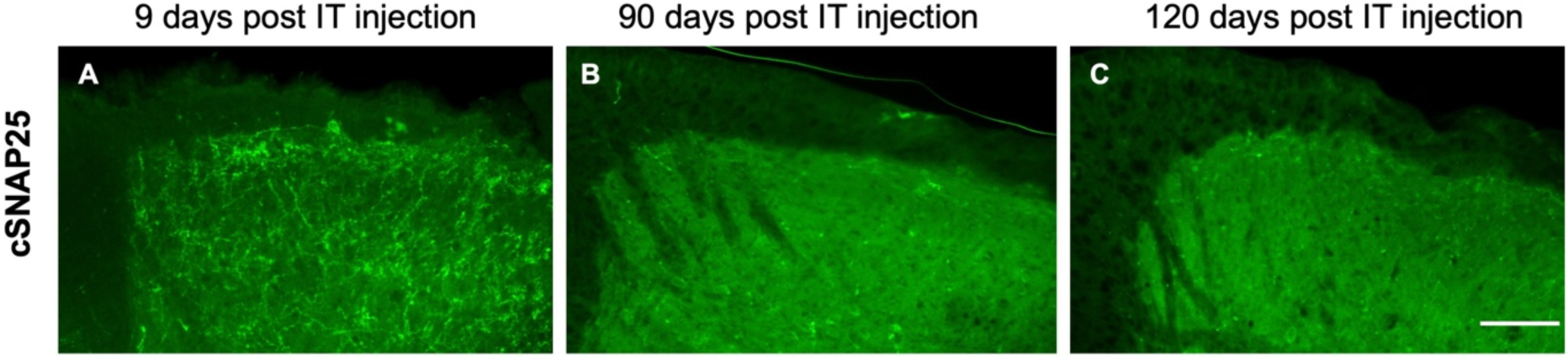
cSNAP immunoreactivity is detectable at 9d post intrathecal SP-BOT but is almost completely absent by 90d and 120d. Confocal immunofluorescence images showing cSNAP25 immunoreactivity (green) in the spinal cord dorsal horn at (A) 9 days, (B) 90 days, and (C) 120 days post-intrathecal (IT) injection of SP-BOT. Scale bar=200μm

**Figure S3:**
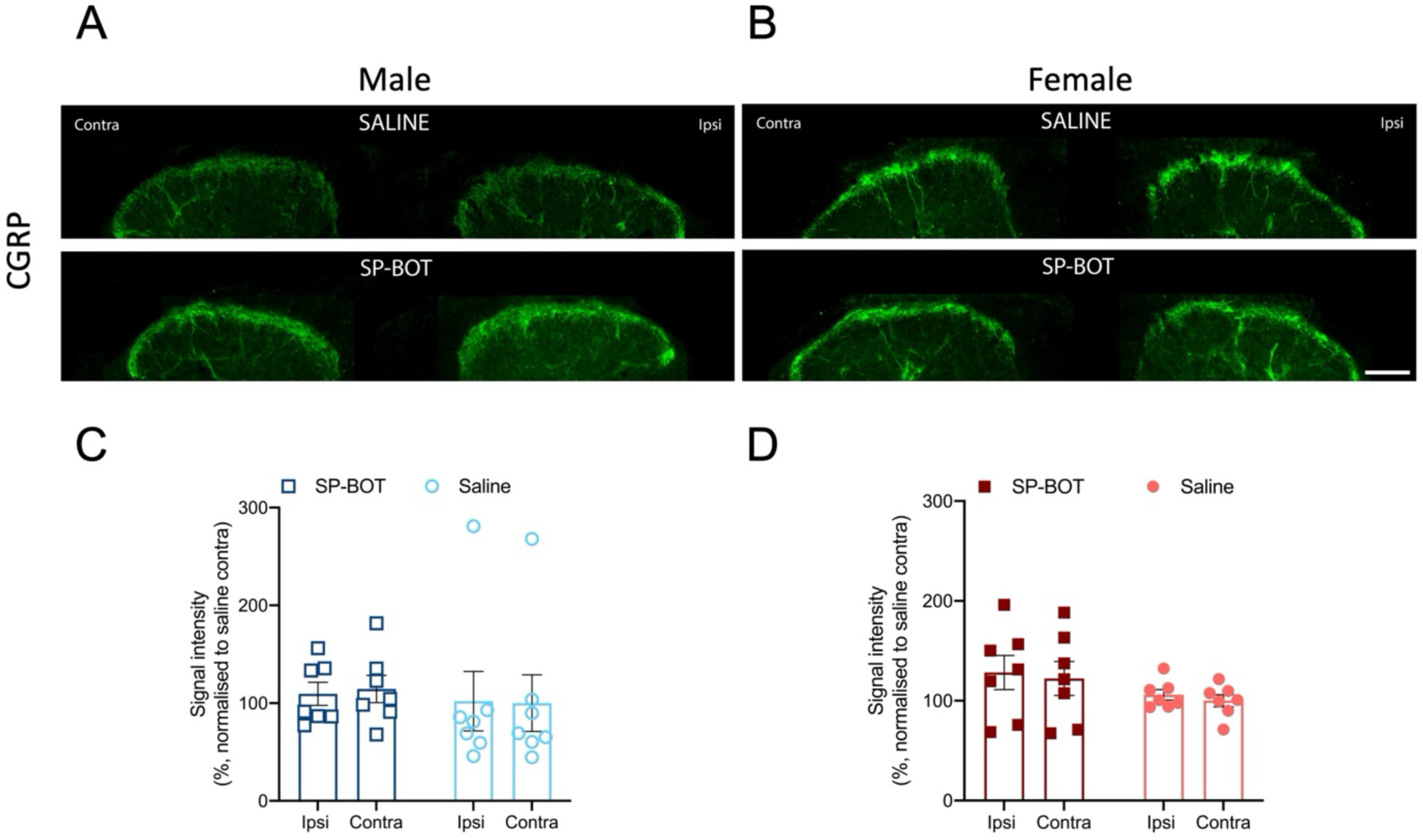
No changes in CGRP levels in mice after CFA injection and SP-BOT or saline intrathecal treatment. **(A-B)** Representative images of CGRP immunoreactivity in the superficial dorsal horn of male and female mice 130 days after injection of CFA in the left ankle joint and 123 days after intrathecal injection of SP-BOT or saline vehicle. Green, CGRP. Scale bar, 100μm. **(C-D)** Quantification of CGRP fluorescence in the ipsilateral and contralateral superficial dorsal horn of male and female mice 130 days after injection of CFA in the left ankle joint and 123 days after intrathecal injection of SP-BOT or saline vehicle. All data were normalised to contralateral saline-treated mice (n=7 per group). Bars represent mean ± SEM on the average fluorescent intensity.

**Figure S4:**
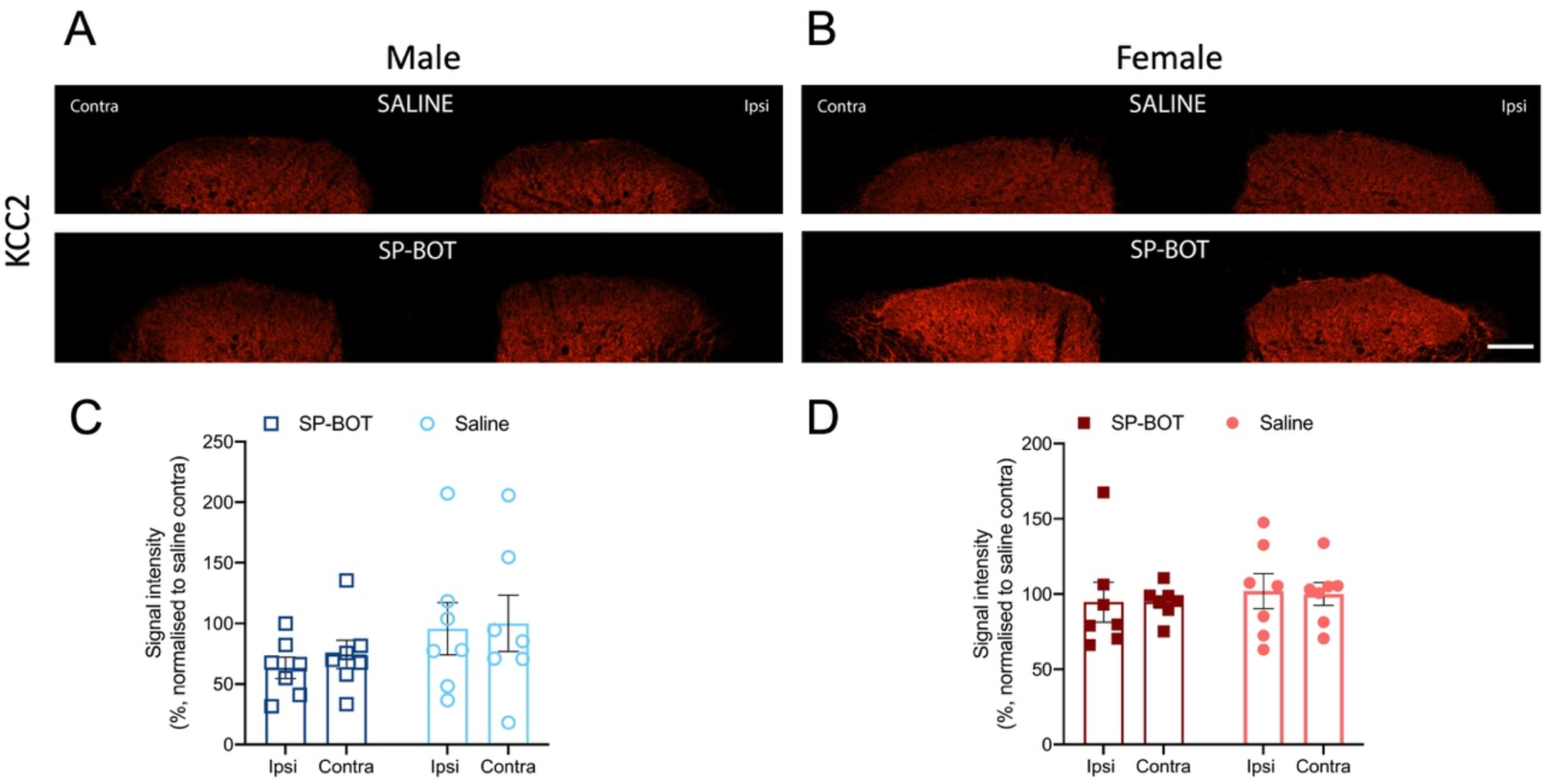
Intrathecal injection of SP-BOT did not reduce KCC2 immunoreactivity in the superficial dorsal horn. **(A-B)** Representative images of KKC2 immunoreactivity in the superficial dorsal horn of male and female mice 130 days after injection of CFA in the left ankle joint and 123 days after intrathecal injection of SP-BOT or saline vehicle. Red, KKC2. Scale bar, 100μm. **(C-D)** quantification of KCC2 fluorescence in the ipsilateral and contralateral superficial dorsal horn of male and female mice 130 days after CFA injection into the left ankle joint and 123 days after intrathecal injection of SP-BOT or saline vehicle. All data were normalised to contralateral saline-treated mice (n=7 per group). Bars represent mean ± SEM on the average fluorescent intensity.

**Figure S5:**
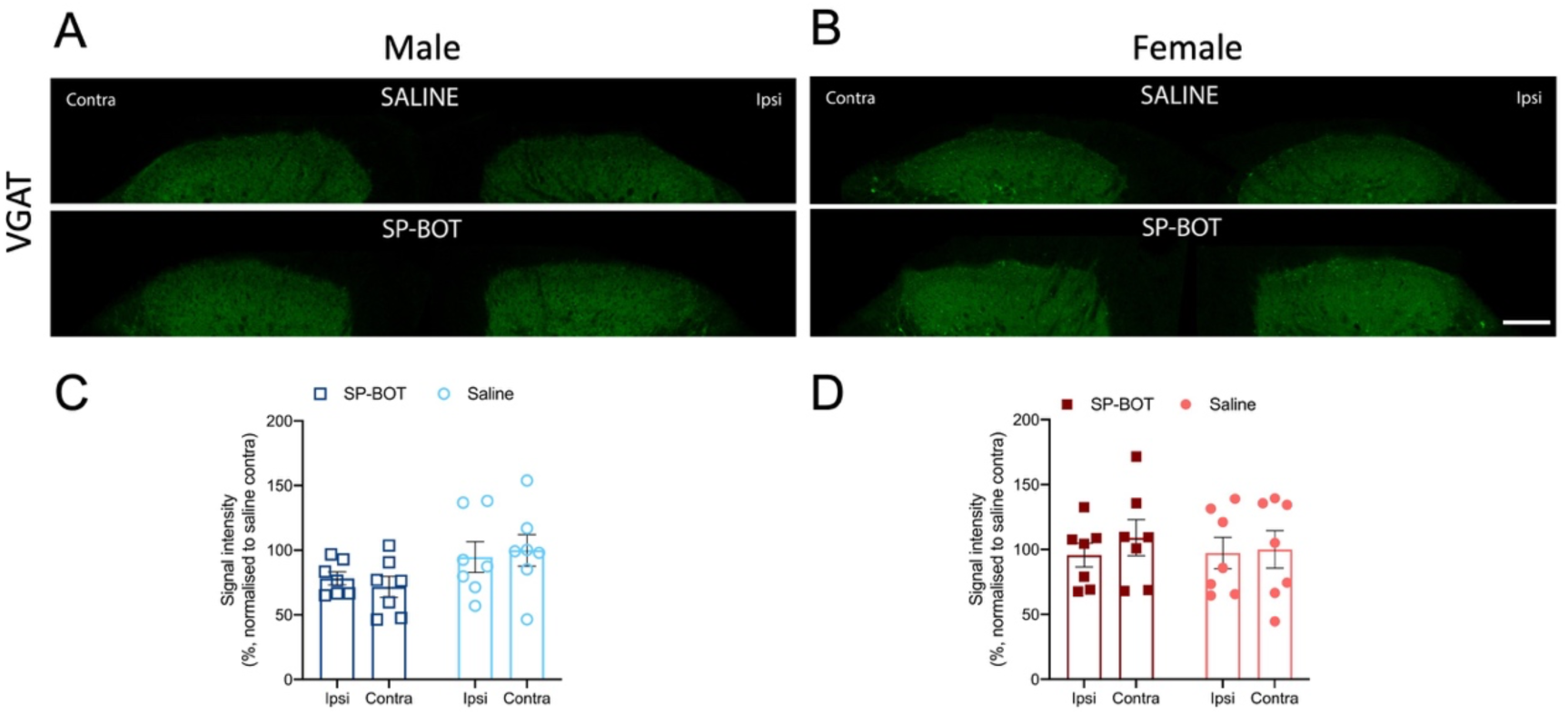
Intrathecal injection of SP-BOT does not modulate levels of VGAT immunoreactivity in the superficial dorsal horn. **(A-B)** Representative images of VGAT immunoreactivity in the superficial dorsal horn of male and female mice 130 days after injection of CFA in the left ankle joint and 123 days after intrathecal injection of SP-BOT or saline vehicle. Green, VGAT. Scale bar, 100μm. **(C-D)** Quantification of VGAT fluorescence in the ipsilateral and contralateral superficial dorsal horn of male and female mice 130 days after CFA injection and 123 days after intrathecal injection of SP-BOT or saline vehicle. All data were normalised to contralateral saline-treated mice (n=7 per group). Bars represent mean ± SEM on the average fluorescent intensity.

## References

[1] Angeby Moller K, Aulin C, Baharpoor A, Svensson CI. Pain behaviour assessments by gait and weight bearing in surgically induced osteoarthritis and inflammatory arthritis. Physiol Behav 2020;225:113079.

[2] Antipova V, Hawlitschka A, Mix E, Schmitt O, Drager D, Benecke R, Wree A. Behavioral and structural effects of unilateral intrastriatal injections of botulinum neurotoxin a in the rat model of Parkinson’s disease. J Neurosci Res 2013;91(6):838–847.

[3] Arsenault J, Ferrari E, Niranjan D, Cuijpers SA, Gu C, Vallis Y, O’Brien J, Davletov B. Stapling of the botulinum type A protease to growth factors and neuropeptides allows selective targeting of neuroendocrine cells. J Neurochem 2013;126(2):223–233.

[4] Berke MS, Hansen CP, Kromann S, Colding-Jorgensen P, Kalliokoski O, Jensen HE, Sorensen DB, Hau J, Abelson KSP, Hestehave S. Refining the adjuvant-induced rat model of monoarthritis by optimizing the induction volume and injection site. Sci Rep 2025;15(1):40281.

[5] Billante CR, Zealear DL, Billante M, Reyes JH, Sant’Anna G, Rodriguez R, Stone RE, Jr. Comparison of neuromuscular blockade and recovery with botulinum toxins A and F. Muscle Nerve 2002;26(3):395–403.

[6] Brown DC, Agnello K. Intrathecal substance P-saporin in the dog: efficacy in bone cancer pain. Anesthesiology 2013;119(5):1178–1185.

[7] Cameron D, Polgar E, Gutierrez-Mecinas M, Gomez-Lima M, Watanabe M, Todd AJ. The organisation of spinoparabrachial neurons in the mouse. Pain 2015;156(10):2061–2071.

[8] Chaplan SR, Bach FW, Pogrel JW, Chung JM, Yaksh TL. Quantitative assessment of tactile allodynia in the rat paw. J Neurosci Methods 1994;53(1):55–63.

[9] Choi S, Hachisuka J, Brett MA, Magee AR, Omori Y, Iqbal NU, Zhang D, DeLisle MM, Wolfson RL, Bai L, Santiago C, Gong S, Goulding M, Heintz N, Koerber HR, Ross SE, Ginty DD. Parallel ascending spinal pathways for affective touch and pain. Nature 2020;587(7833):258–263.

[10] Corder G, Doolen S, Donahue RR, Winter MK, Jutras BL, He Y, Hu X, Wieskopf JS, Mogil JS, Storm DR, Wang ZJ, McCarson KE, Taylor BK. Constitutive mu-opioid receptor activity leads to long-term endogenous analgesia and dependence. Science 2013;341(6152):1394–1399.

[11] Corder G, Tawfik VL, Wang D, Sypek EI, Low SA, Dickinson JR, Sotoudeh C, Clark JD, Barres BA, Bohlen CJ, Scherrer G. Loss of mu opioid receptor signaling in nociceptors, but not microglia, abrogates morphine tolerance without disrupting analgesia. Nat Med 2017;23(2):164–173.

[12] Darios F, Niranjan D, Ferrari E, Zhang F, Soloviev M, Rummel A, Bigalke H, Suckling J, Ushkaryov Y, Naumenko N, Shakirzyanova A, Giniatullin R, Maywood E, Hastings M, Binz T, Davletov B. SNARE tagging allows stepwise assembly of a multimodular medicinal toxin. Proc Natl Acad Sci U S A 2010;107(42):18197–18201.

[13] Fairbanks CA. Spinal delivery of analgesics in experimental models of pain and analgesia. Adv Drug Deliv Rev 2003;55(8):1007–1041.

[14] Fatt MP, Zhang MD, Kupari J, Altinkok M, Yang Y, Hu Y, Svenningsson P, Ernfors P. Morphine-responsive neurons that regulate mechanical antinociception. Science 2024;385(6712):eado6593.

[15] Ferdousi M, Finn DP. Stress-induced modulation of pain: Role of the endogenous opioid system. Prog Brain Res 2018;239:121–177.

[16] Ferrari E, Gu C, Niranjan D, Restani L, Rasetti-Escargueil C, Obara I, Geranton SM, Arsenault J, Goetze TA, Harper CB, Nguyen TH, Maywood E, O’Brien J, Schiavo G, Wheeler DW, Meunier FA, Hastings M, Edwardson JM, Sesardic D, Caleo M, Hunt SP, Davletov B. Synthetic self-assembling clostridial chimera for modulation of sensory functions. Bioconjug Chem 2013;24(10):1750–1759.

[17] Finnerup NB, Attal N, Haroutounian S, McNicol E, Baron R, Dworkin RH, Gilron I, Haanpaa M, Hansson P, Jensen TS, Kamerman PR, Lund K, Moore A, Raja SN, Rice AS, Rowbotham M, Sena E, Siddall P, Smith BH, Wallace M. Pharmacotherapy for neuropathic pain in adults: a systematic review and meta-analysis. Lancet Neurol 2015;14(2):162–173.

[18] Francois A, Low SA, Sypek EI, Christensen AJ, Sotoudeh C, Beier KT, Ramakrishnan C, Ritola KD, Sharif-Naeini R, Deisseroth K, Delp SL, Malenka RC, Luo L, Hantman AW, Scherrer G. A Brainstem-Spinal Cord Inhibitory Circuit for Mechanical Pain Modulation by GABA and Enkephalins. Neuron 2017;93(4):822–839 e826.

[19] Gerum M, Simonin F. Behavioral characterization, potential clinical relevance and mechanisms of latent pain sensitization. Pharmacol Ther 2022;233:108032.

[20] Haywood AR, Hathway GJ, Chapman V. Differential contributions of peripheral and central mechanisms to pain in a rodent model of osteoarthritis. Sci Rep 2018;8(1):7122.

[21] Hestehave S, Florea R, Fedorec AJH, Jevic M, Mercy L, Wright A, Morgan OB, Brown LA, Peirson SN, Geranton SM. Differences in multidimensional phenotype of 2 joint pain models link early weight-bearing deficit to late depressive-like behavior in male mice. Pain Rep 2024;9(6):e1213.

[22] Hestehave S, Florea R, Singleton S, Fedorec AJH, Caxaria S, Morgan OB, Kopp KT, Brown LA, Heymann T, Sikandar S, Hausch F, Peirson SN, Geranton SM. Analgesia through FKBP51 inhibition at disease onset confers lasting relief from sensory and emotional chronic pain symptoms. Proc Natl Acad Sci U S A 2025;122(44):e2517405122.

[23] Khasabov SG, Rogers SD, Ghilardi JR, Peters CM, Mantyh PW, Simone DA. Spinal neurons that possess the substance P receptor are required for the development of central sensitization. J Neurosci 2002;22(20):9086–9098.

[24] Leese C, Christmas C, Meszaros J, Ward S, Maiaru M, Hunt SP, Davletov B. New botulinum neurotoxin constructs for treatment of chronic pain. Life Sci Alliance 2023;6(6).

[25] Locke S, Yousefpour N, Mannarino M, Xing S, Yashmin F, Bourassa V, Ribeiro-da-Silva A. Peripheral and central nervous system alterations in a rat model of inflammatory arthritis. Pain 2020;161(7):1483–1496.

[26] Maiaru M, Leese C, Certo M, Echeverria-Altuna I, Mangione AS, Arsenault J, Davletov B, Hunt SP. Selective neuronal silencing using synthetic botulinum molecules alleviates chronic pain in mice. Sci Transl Med 2018;10(450).

[27] Maiaru M, Leese C, Silva-Hucha S, Fontana-Giusti S, Tait L, Tamagnini F, Davletov B, Hunt SP. Substance P-Botulinum Mediates Long-term Silencing of Pain Pathways that can be Re-instated with a Second Injection of the Construct in Mice. J Pain 2024;25(6):104466.

[28] Maiaru M, Tochiki KK, Cox MB, Annan LV, Bell CG, Feng X, Hausch F, Geranton SM. The stress regulator FKBP51 drives chronic pain by modulating spinal glucocorticoid signaling. Sci Transl Med 2016;8(325):325ra319.

[29] Mantyh PW, Rogers SD, Honore P, Allen BJ, Ghilardi JR, Li J, Daughters RS, Lappi DA, Wiley RG, Simone DA. Inhibition of hyperalgesia by ablation of lamina I spinal neurons expressing the substance P receptor. Science 1997;278(5336):275–279.

[30] Mills C, Leblond D, Joshi S, Zhu C, Hsieh G, Jacobson P, Meyer M, Decker M. Estimating efficacy and drug ED50’s using von Frey thresholds: impact of weber’s law and log transformation. J Pain 2012;13(6):519–523.

[31] Mogil JS, Parisien M, Esfahani SJ, Diatchenko L. Sex differences in mechanisms of pain hypersensitivity. Neurosci Biobehav Rev 2024;163:105749.

[32] Nichols ML, Allen BJ, Rogers SD, Ghilardi JR, Honore P, Luger NM, Finke MP, Li J, Lappi DA, Simone DA, Mantyh PW. Transmission of chronic nociception by spinal neurons expressing the substance P receptor. Science 1999;286(5444):1558–1561.

[33] Ninkovic M, Hunt SP, Kelly JS. Effect of dorsal rhizotomy on the autoradiographic distribution of opiate and neurotensin receptors and neurotensin-like immunoreactivity within the rat spinal cord. Brain Res 1981;230(1-2):111–119.

[34] Ohashi Y, Uchida K, Fukushima K, Inoue G, Takaso M. Mechanisms of Peripheral and Central Sensitization in Osteoarthritis Pain. Cureus 2023;15(2):e35331.

[35] Pavone F, Luvisetto S. Botulinum neurotoxin for pain management: insights from animal models. Toxins (Basel) 2010;2(12):2890–2913.

[36] Presto P, Mazzitelli M, Junell R, Griffin Z, Neugebauer V. Sex differences in pain along the neuraxis. Neuropharmacology 2022;210:109030.

[37] Rivat C, Laboureyras E, Laulin JP, Le Roy C, Richebe P, Simonnet G. Non-nociceptive environmental stress induces hyperalgesia, not analgesia, in pain and opioid-experienced rats. Neuropsychopharmacology 2007;32(10):2217–2228.

[38] Ruscheweyh R, Athwal B, Gryglas-Dworak A, Frattale I, Latysheva N, Ornello R, Pozo-Rosich P, Sacco S, Torres Ferrus M, Stark CD. Wear-Off of OnabotulinumtoxinA Effect Over the Treatment Interval in Chronic Migraine: A Retrospective Chart Review With Analysis of Headache Diaries. Headache 2020;60(8):1673–1682.

[39] Segal NA, Nilges JM, Oo WM. Sex differences in osteoarthritis prevalence, pain perception, physical function and therapeutics. Osteoarthritis Cartilage 2024;32(9):1045–1053.

[40] Severino A, Chen W, Hakimian JK, Kieffer BL, Gaveriaux-Ruff C, Walwyn W, Marvizon JCG. Mu-opioid receptors in nociceptive afferents produce a sustained suppression of hyperalgesia in chronic pain. Pain 2018;159(8):1607–1620.

[41] Sheahan TD, Warwick CA, Fanien LG, Ross SE. The Neurokinin-1 Receptor is Expressed with Gastrin-Releasing Peptide Receptor in Spinal Interneurons and Modulates Itch. J Neurosci 2020;40(46):8816–8830.

[42] Smith AF, Plumb AN, Berardi G, Sluka KA. Sex differences in the transition to chronic pain. J Clin Invest 2025;135(11).

[43] Solway B, Bose SC, Corder G, Donahue RR, Taylor BK. Tonic inhibition of chronic pain by neuropeptide Y. Proc Natl Acad Sci U S A 2011;108(17):7224–7229.

[44] Suzuki R, Morcuende S, Webber M, Hunt SP, Dickenson AH. Superficial NK1-expressing neurons control spinal excitability through activation of descending pathways. Nat Neurosci 2002;5(12):1319–1326.

[45] Syx D, Tran PB, Miller RE, Malfait AM. Peripheral Mechanisms Contributing to Osteoarthritis Pain. Curr Rheumatol Rep 2018;20(2):9.

[46] Todd AJ, Hughes DI, Polgar E, Nagy GG, Mackie M, Ottersen OP, Maxwell DJ. The expression of vesicular glutamate transporters VGLUT1 and VGLUT2 in neurochemically defined axonal populations in the rat spinal cord with emphasis on the dorsal horn. Eur J Neurosci 2003;17(1):13–27.

[47] Vierck CJ, Jr., Kline RH, Wiley RG. Intrathecal substance p-saporin attenuates operant escape from nociceptive thermal stimuli. Neuroscience 2003;119(1):223–232.

[48] Vincent TL. Peripheral pain mechanisms in osteoarthritis. Pain 2020;161 Suppl 1(1):S138–S146.

[49] Vrinten DH, Hamers FF. ’CatWalk’ automated quantitative gait analysis as a novel method to assess mechanical allodynia in the rat; a comparison with von Frey testing. Pain 2003;102(1-2):203–209.

[50] Walwyn WM, Chen W, Kim H, Minasyan A, Ennes HS, McRoberts JA, Marvizon JC. Sustained Suppression of Hyperalgesia during Latent Sensitization by mu-, delta-, and kappa-opioid receptors and alpha2A Adrenergic Receptors: Role of Constitutive Activity. J Neurosci 2016;36(1):204–221.

[51] Weibel R, Reiss D, Karchewski L, Gardon O, Matifas A, Filliol D, Becker JA, Wood JN, Kieffer BL, Gaveriaux-Ruff C. Mu opioid receptors on primary afferent nav1.8 neurons contribute to opiate-induced analgesia: insight from conditional knockout mice. PLoS One 2013;8(9):e74706.

[52] Zhang Y, Zhao S, Rodriguez E, Takatoh J, Han BX, Zhou X, Wang F. Identifying local and descending inputs for primary sensory neurons. J Clin Invest 2015;125(10):3782–3794.

